# The yellow-in-the-dark *chIL* Chlamydomonas mutant as a model for time-resolved chloroplast biogenesis and physiological responses to lincomycin

**DOI:** 10.64898/2025.12.12.693793

**Authors:** M. Águila Ruiz-Sola, Mariano A. De Silvio, Mario Becerra-Tinoco, Andrea M. Quintero-Moreno, José L. Crespo, Elena Monte

## Abstract

Understanding chloroplast biogenesis is key given the importance of photosynthesis and chloroplast metabolism. Chloroplast assembly has been extensively studied in angiosperms, where the main hallmarks have been established, including the need for nuclear and plastid genome coordination through anterograde and retrograde signals. However, recent works in other photosynthetic lineages have revealed both divergent and conserved regulators. These differences likely represent adaptive strategies that emerged during the evolution of the green lineage, and highlight the need for evolutionary studies to fully elucidate the mechanisms controlling chloroplast biogenesis, a light-promoted process. Here, we characterize the yellow-in-the-dark *chIL* mutant of *Chlamydomonas reinhardtii*, which carries a mutation in the CHLL subunit of the dark-operative POR enzyme. Dark-adapted *chIL* cells cannot synthesize chlorophylls and display a de-differentiated chloroplast, and our results show that upon exposure to light, chloroplast biogenesis is initiated very rapidly based on starch reserves. We demonstrate that accumulation of pigments, photosynthesis-related proteins, and functional chloroplast activity is detected within 1-3 hours of illumination. Additionally, we show that lincomycin, an inhibitor of plastid translation, specifically affects chloroplast physiology, and blocks light-induced chloroplast biogenesis and cell growth, establishing lincomycin as a valuable tool for investigating chloroplast physiology and plastid-to-nucleus signals in *Chlamydomonas*. Together, this study introduces the *chIL* mutant as a powerful model system to dissect light-induced, time-resolved chloroplast biogenesis, as well as the responses and signaling triggered by lincomycin. By promoting broader adoption of this model, we aim to help inform strategies to enhance photosynthesis and stress tolerance, and optimize bioproduct production.

## Introduction

Chloroplasts house photosynthesis and function as metabolic hubs necessary for pigment biosynthesis and production of vital metabolites (Rolland et al., 2018). Understanding chloroplast biogenesis is key to establish the basis of photosynthetic and metabolic competence, essential for adaptation to changing environments and global primary production.

Chloroplast biogenesis has been extensively studied in angiosperms. Dark-grown etiolated seedlings accumulate etioplasts featuring a prolamellar body (PLB), which harbours the chlorophyll precursor protochlorophyllide (Pchlide) and the nuclear-encoded, light-dependent enzyme Pchlide oxidoreductase (LPOR) that controls the reduction of Pchlide to chlorophyllide (Chlide) in the light (Floris and Kühlbrandt, 2021). Consequently, etiolated seedlings lack chlorophylls and exhibit a yellowish colour characteristic of carotenoids. Upon exposure to light, etioplasts develop into green chloroplasts and seedlings undergo de-etiolation to promote optimal photosynthetic performance (Gommers and Monte, 2018).

In contrast to flowering plants, most of the remaining major groups of the green lineage retain the capacity to produce chlorophyll in the dark thanks to a light-independent Dark-operative Pchlide Oxidoreductase (DPOR) (Vedalankar and Tripathy, 2019). DPOR is plastid-encoded and consists of three subunits (CHLN, CHLB, and CHLL), which catalyze the light-independent reduction of Pchlide to Chlide (Armstrong, 1998). DPOR activity has been lost entirely in angiosperms and partially in gymnosperms and some algae (Solymosi and Schoefs, 2010; Hunsperger et al., 2015). Importantly, chlorophyte algae, including the model organism *Chlamydomonas reinhardtii*, retains high DPOR prevalence.

Numerous studies in angiosperms have defined the hallmarks of chloroplast biogenesis (Jarvis and López-Juez, 2013; Pipitone et al., 2021), which include the coordinated synthesis and import of the chloroplast protein machinery encoded in the nuclear genome, with the massive production of lipids and chlorophylls in the chloroplast to drive the assembly of the photosynthetic complexes and thylakoid membranes. Light is essential for activating the transcriptomic reprogramming necessary for chloroplast biogenesis, and many of the regulators have been elucidated (Frangedakis et al., 2024). Recent works in other photosynthetic lineages have revealed both divergent and conserved regulators, including some previously overlooked in Arabidopsis (Nishiyama et al., 2018; Frangedakis et al., 2024). Such differences likely reflect adaptive strategies that emerged throughout the evolution of the green lineage, highlighting the need for comparative studies to fully uncover the mechanisms that control light-regulated chloroplast biogenesis.

The unicellular green alga *Chlamydomonas reinhardtii*, featuring a single chloroplast with thylakoid architecture and composition similar to higher plants, provides a powerful system to explore the evolutionary foundations of chloroplast biogenesis (Ostermeier et al., 2024). Chlamydomonas can grow non-photosynthetically in the dark in acetate-containing media while maintaining a green and functional chloroplast due to DPOR activity (Harris, 2001). Importantly, a yellow-in-the-dark mutant *y-1*, with defects in the accumulation of the DPOR subunit CHLL, was used to pioneer investigations into light-regulated chloroplast biogenesis (SAGER and PALADE, 1954; Ohad et al., 1967a; Cahoon and Timko, 2000). Similar to etiolated seedlings, *y-1* are yellowish with undifferentiated plastids that accumulate Pchlide and carotenoids in darkness and fully develop into functional chloroplasts upon illumination (Ohad et al., 1967b; Malnoë et al., 1988; White and Hoober, 1994). Unfortunately, the genetic basis of *y-1* remains unknown, hindering its use as a model. This was addressed by targeted chloroplast mutations in the DPOR CHLL subunit, that generated the yellow-in-the-dark *chlL* mutant (Suzuki and Bauer, 1992). However, chloroplast biogenesis in this *chlL* mutant remains to be characterized.

The antibiotic lincomycin inhibits protein synthesis in the chloroplast and has been extensively used in plants to study chloroplast development and the coordination of nuclear and plastid genomes (León et al., 2025). In contrast, the use of lincomycin to study chloroplast biogenesis and signalling in Chlamydomonas has not been explored.

Here, we describe a detailed methodology to establish the *chlL* mutant as a model system to study time-resolved light-induced chloroplast biogenesis. We provide a comprehensive physiological and biochemical analysis of chloroplast biogenesis in *chlL* from a dark undifferentiated plastid to a fully functional chloroplast. Furthermore, we show that the antibiotic lincomycin is a valuable tool for investigating the process of chloroplast biogenesis in Chlamydomonas.

## Results

### Lincomycin impacts Chlamydomonas growth and physiology

Wild-type strains CC-124 and CC-3269 were grown on the acetate-containing TAP and minimum medium HSM plates in continuous light (TAP Light and TP Light) or darkness (TAP Dark) in the presence or absence of lincomycin. Growth in TP Light was slower compared to TAP Light as expected, whereas growth in TAP Dark took longer, as shown in the serial spot dilutions in Figure 1A depicting cells grown for 6, 5 and 12 days, respectively. In light conditions, presence of lincomycin considerably affected growth, whereas in darkness lincomycin did not appear to have a visible impact (Figure 1A). CC-124 grown under the same conditions in liquid medium (Figure1B) showed exponential growth after inoculation (day 0) in TAP Light from day 1 to day 3, when it reached a plateau. In TP Light, and even more in TAP Dark, growth was slower (Figure 1B), mirroring the dynamics on plates (Figure 1A). Likewise, addition of lincomycin prevented growth only in light conditions (Figure 1B). Corresponding growth rates showed an increase between days 0 and 2 in TAP Light and Dark, and between days 0 and 3 in HSM Light, with a subsequent drop (Figure 1C). Again, lincomycin-treated samples showed reduced growth rate only in light samples (Figure 1C). Together, our results show that lincomycin impairs the growth of WT Chlamydomonas strains in the light in both acetate-containing and minimal medium, whereas no significant effect was detected under dark conditions.

**Figure 1.**
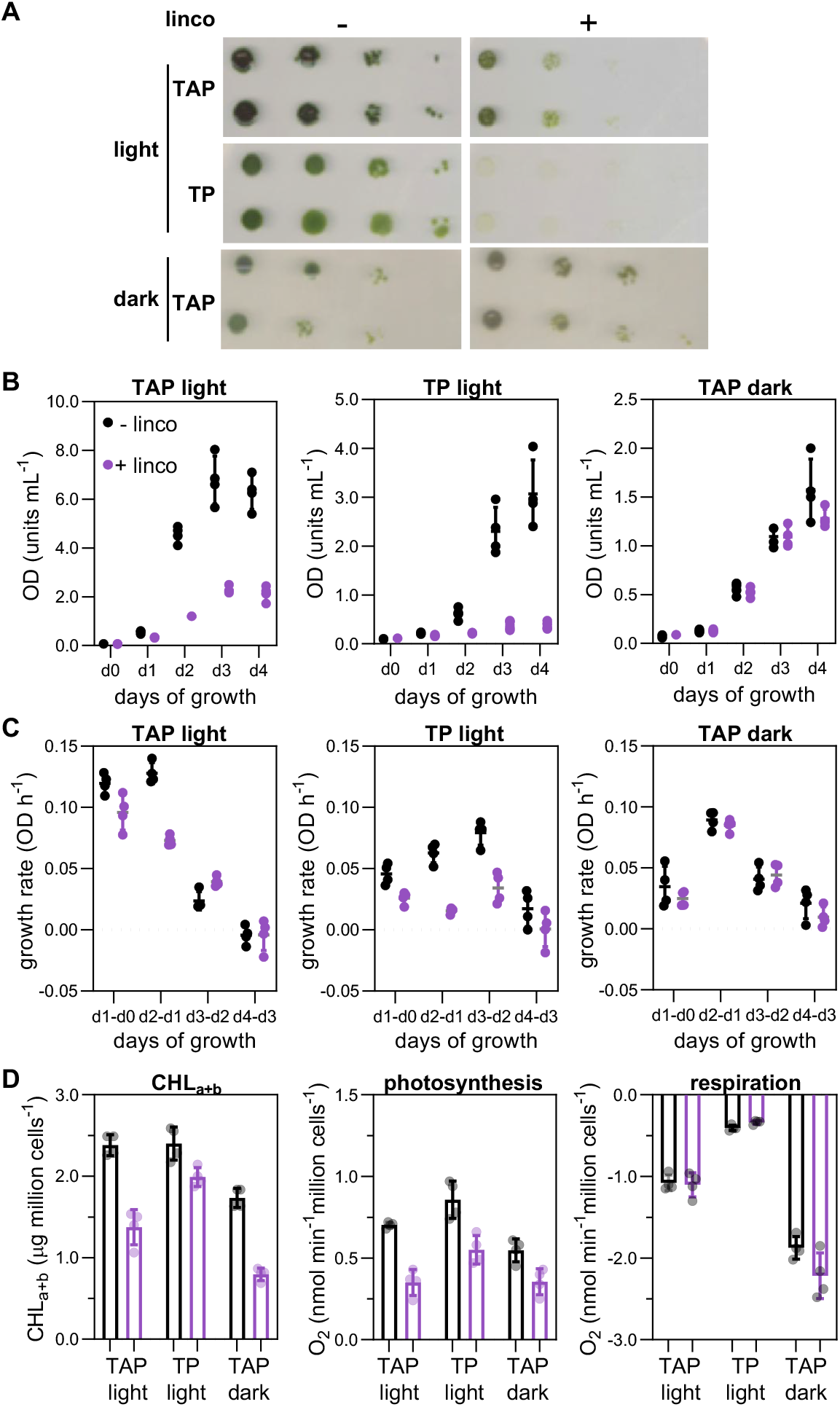
Lincomycin affects Chlamydomonas growth and physiology. A, Growth of Chlamydomonas strains CC-124 and CC-3269 under continuous light in solid TAP (5 days) and TP (6 days) medium, or under continuous darkness on TAP (12 days). Cells were subjected to 10-fold serial dilutions and spotted into plates with or without lincomycin 0.5 mM (Linco). B, Growth curves of CC-124 grown in liquid medium as in A, for 0, 1, 2, 3, and 4 days after inoculation, expressed as O.D._750_ units/mL. C, Growth rate (expressed as O.D. _750_ units/hour) of cultures grown in B. D, Total chlorophylls, and photosynthesis and respiration rates of CC-124 cells after 1 day of growth as in B.

We next complemented the growth data with physiological measurements at day 1. Chlorophyll content was similar in TAP Light and TP Light, but reduced in TAP Dark, and lincomycin treatment triggered a decrease in all three growth conditions (Figure 1D). Measurement of photosynthetic oxygen evolution indicated that TAP Light and TP Light cells exhibited similar photosynthetic capacity (Figure 1D), while TAP Dark cells had reduced levels. Interestingly, in all three conditions, photosynthesis was affected by lincomycin (Figure 1D). Finally, mitochondrial respiration measured by assessing oxygen consumption in the dark showed some differences among light treatments, likely reflecting distinct mitochondrial activity during mixotrophic, autotrophic and heterotrophic growth, but remarkably no differences were observed between lincomycin-treated and non-treated samples in each condition. Our results indicate that lincomycin can impact chlorophyll content and photosynthesis in cells grown in both light and dark, and this effect on chloroplast physiology appears to be specific as lincomycin does not affect respiration significantly.

### Chloroplast structure and pigment profile of the yellow-in-the-dark *chlL* mutant under different light regimes

We next established a detailed protocol to characterize the physiology and subcellular features of the *chlL* mutant grown under our conditions. *chlL* growth in TAP Light, TP Light and TAP Dark showed similar growth patterns compared to the WT control CC-3269 (Figure 2A), indicating that *chlL* cells can grow and divide in both light and dark conditions. *chlL* grown on TAP Light plates was used to inoculate a liquid TAP culture and placed in continuous darkness. Under these conditions, the *chlL* mutant turned yellowish after 5 days (Figure 2B), as expected due to its DPOR deficiency, which prevents chlorophyll accumulation in the dark (Suzuki and Bauer, 1992). However, in agreement with LPOR activation upon light illumination, dark-grown *chlL* cultures transferred to TP and light for 48h were green and undistinguishable from continuous light-grown *chlL* (Figure 2B).

**Figure 2.**
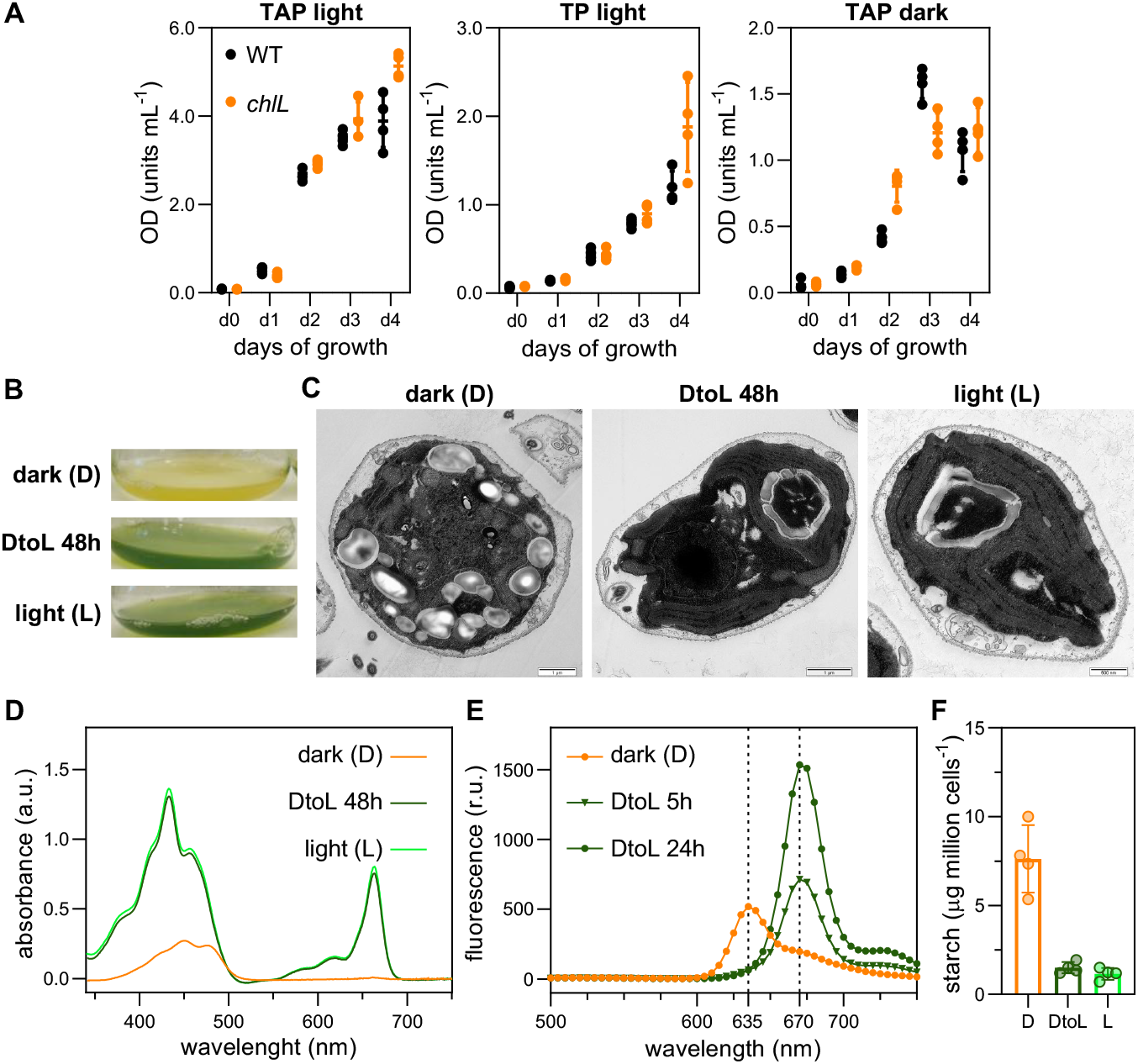
Chloroplast structure and pigment profile of the *chlL* mutant. A, Growth of Chlamydomonas strains CC-3269 and *chlL* in liquid TAP or TP medium in continuous light, or under continuous darkness in TAP for 0, 1, 2, 3, and 4 days after inoculation, expressed as O.D._750_ units/mL. B, Appearance of liquid cultures of *chIL* cells grown for 7 days in TAP medium in the dark (top), for 7 days under continuous light in TP (bottom), or grown in the dark for 5 days in TAP and then transferred to light in TP for 48h (middle). C, Transmission electron microscopy (TEM) images of *chIL* cells grown as in B. D, Absorbance spectra of *chIL* cells grown as in B, with values expressed as arbitrary units (a.u.). E, Fluorescence spectrum after excitation at 440 nm of dark-grown *chIL* cells (0h) transferred to light for 5h and 24h, expressed as relative units (r.u.). F, Starch quantification of *chIL* cells grown as in B, expressed as µg of starch per million cells.

At the subcellular level, dark-grown *chlL* cells exhibited a partially developed chloroplast that lacked thylakoid membranes and grana stacks (Figure 2C), similar to the *y-1* mutant (Ohad et al., 1967a). Dark-grown *chlL* cells transferred to light for 48h showed fully developed cup-shaped chloroplasts with thylakoid membranes stacked in grana, like *chlL* cells grown under continuous light (Figure 2C). Photosynthetic pigment accumulation in all three conditions was in accordance to these phenotypes: dark-grown *chlL* mutant showed accumulation of carotenoids but no chlorophylls (Figure 2D), whereas dark-grown *chlL* cells transferred to light for 48h and light-grown *chlL* cells accumulated carotenoids and chlorophylls to similar amounts (Figure 2D). During the dark-to-light conditions, the chlorophyll precursor protochlorophyllide is rapidly transformed into chlorophyllide by the LPOR enzyme (Figure 2E). Together, these results indicate that dark-grown yellow *chlL* cells transferred to light accumulate chlorophyll and develop a structured chloroplast similar to *chlL* cells grown under continuous light. Interestingly, we observed presence of starch granules in TAP Dark that were absent after 48h of light, presumably consumed to fuel chloroplast biogenesis (Figures 2C and 2F).

### Dark-grown yellow *chlL* mutant rapidly builds a functional chloroplast when exposed to light, a process impaired in the presence of lincomycin

To establish the timing and physiology of chloroplast biogenesis in *chIL*, cells were first grown in continuous darkness until yellow and were then transferred to continuous light (time point= 0h) for 48h, sampling after 1, 3, 6, 9, 12, 24, and 48h. A separate culture of *chIL* cells was grown under continuous light as reference control. Throughout the dark-to-light time-course, cells looked healthy and greening was progressive and visually apparent after 3-6h (Figures 3A and 3B), with cell division measured as total cell count detectable 9-12h after lights on (Figure 3C). Chlorophyll and carotenoid accumulation (Figures 3D and 3E), protochlorophyllide to chlorophyllide conversion (Figure 3F), and the assembly of functional photosystem (PS) PSII and PSI (as shown by F_v_/F_m_ and P_m_ values respectively) were detected as early as 1-3h after light onset (Figures 3G and 3H, and Supplemental Figure S1). In accordance, photosynthetic activity measured by oxygen production was detected after 3-6h (Figure 3I). Complete chloroplast biogenesis was reached after 48h, when physiological values were similar to control *chIL* cells grown under continuous light (Figures 3D-3I). Overall, physiological values of our *chIL* system were comparable and expanded previous studies using other *yellow-in-the-dark* genotypes (Ohad et al., 1967b; White and Hoober, 1994).

**Figure 3.**
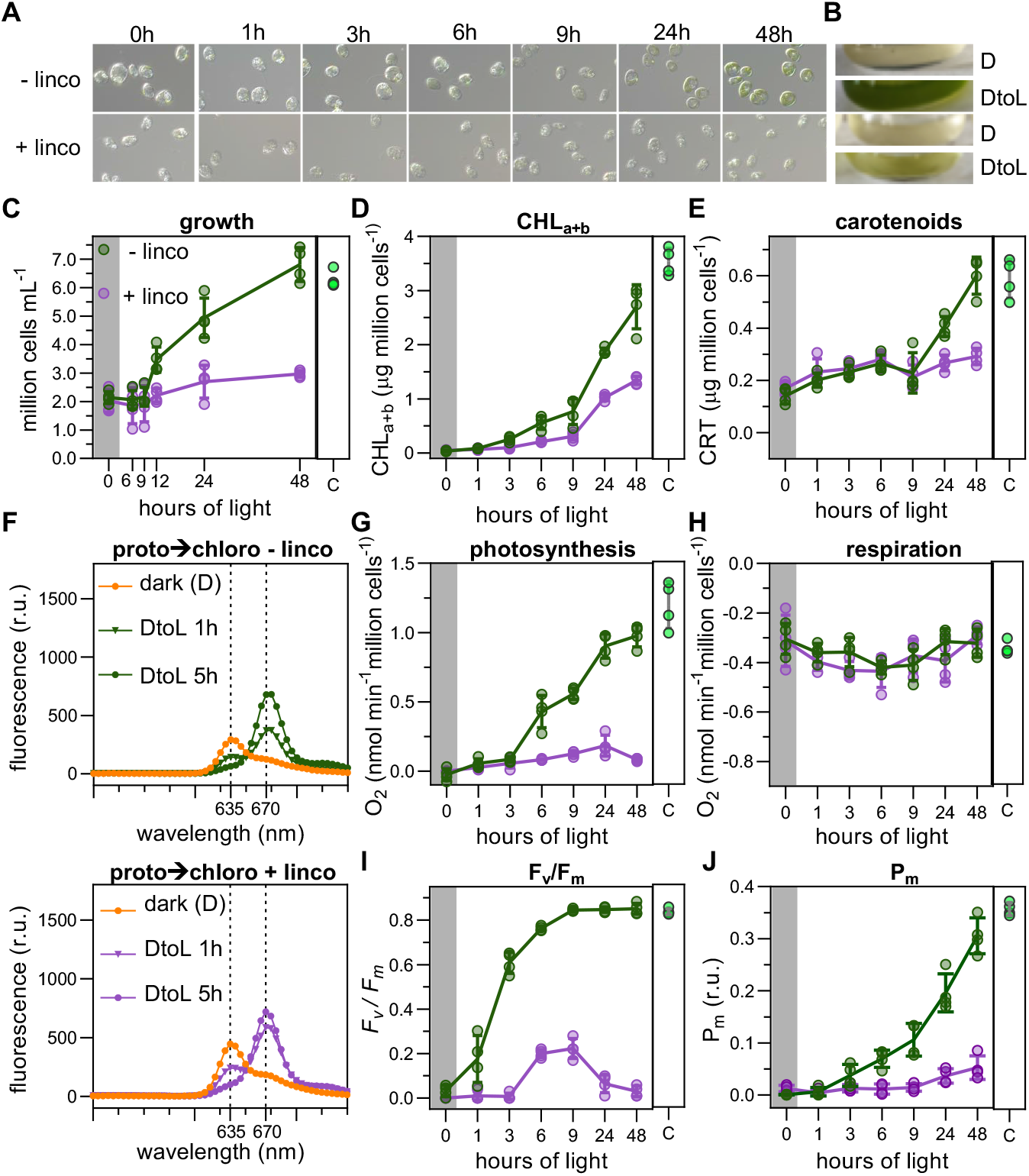
Time-resolved physiology of chloroplast biogenesis in the *chlL* mutant upon transfer to the light in the presence or absence of lincomycin. A, Light micrographs of dark-grown *chIL* cells in liquid cultures exposed to continuous light for the indicated times (0 (Dark), 1, 3, 6, 9, 24, 48h), in the presence (bottom) or absence (top) of lincomycin. B, Appearance of *chIL* ccultures grown as in A. DtoL (Dark to Light) correspond to samples after 48h. C, Growth curves expressed as cell number (million cells/mL). D, Total chlorophyll content expressed in µg per million cells. E, Total carotenoid content expressed in µg per million cells. F, Fluorescence spectrum after excitation at 440 nm of dark-grown *chIL* cells (0h) transferred to light for 1h and 5h, expressed as relative fluoresce units (r.u.). G, F_v_/F_m_ fluorescence values. H, P_m_ values expressed in relative units (r.u.). Green lines correspond to light-treated samples. Purple lines correspond to light and lincomycin-treated samples. The light green sample corresponds to the control (C) *chlL* mutant grown under continuous light.

Addition of lincomycin to the system had a clear impact on greening (Figures 3A and 3B). Although rapid protochlorophyllide-to-chlorophyllide conversion was not affected (Figure 3F), lincomycin impaired the accumulation of chlorophylls and carotenoids, PS assembly and activity, as well as cell division (Figures 3C-E, 3G and 3H, and Supplemental Figure S1). Interestingly, lincomycin increased F0 values, suggesting the presence of free chlorophylls due to impaired photosystem assembly (Supplemental Figure S1). Importantly, O_2_ respiration rate showed similar values in all samples (Figure 3J), indicating that the effect of lincomycin is highly specific to the chloroplast. In short, dark-grown yellow *chIL* cells rapidly assemble a functional chloroplast upon light exposure, a process impaired by the addition of lincomycin.

### Light induces early accumulation of photosynthesis-related proteins during chloroplast biogenesis

To further understand the dynamics of light-induced chloroplast assembly, we next assessed the accumulation of key photosynthesis-related proteins in dark-grown yellow *chIL* cells exposed to light after 0, 1, 3, 6, 9, 12, 24, and 48 h of light treatment, and compared them to *chIL* grown under continuous light as control. Evaluated proteins included the PSII core D1 and CP43 and antenna Lhcbm5, the PSI core PsaA and PsaC, the cytochrome *b6f* subunits PetA and PetB, and the beta-subunit ATPb of the chloroplast ATP synthase. Results are shown in Figure 4 and Supplemental Figure 2, which include immunoblots for two independent replicates (one shown in Figure 4) and their quantification relative to the continuous light control.

**Figure 4.**
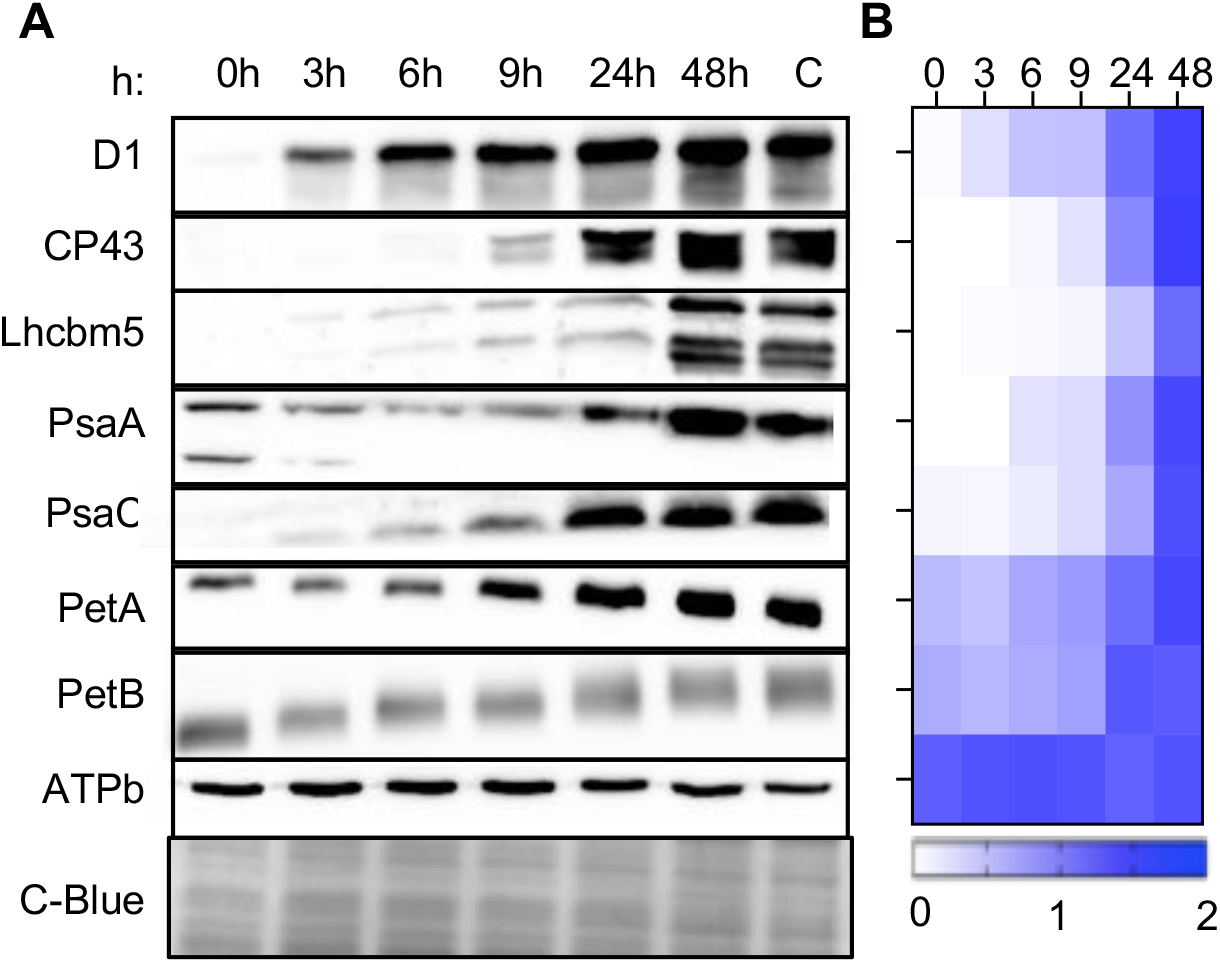
Accumulation of photosynthesis-related proteins during chloroplast biogenesis in *chIL*. A, Immunoblot analysis of the accumulation of PSII (D1 and CP43), PSI (Lhcbm5, PsaA and PsaC), Cyt *b6f* complex (PetA and PetB) and ATP synthase ATPb proteins, in dark-grown *chIL* cells at 0, 1, 3, 6, 9, 12, 24, and 48 h after illumination. A control (C) corresponding to *chIL* cells grown under continuous light is included. Coomassie blue (C-blue) staining served as loading control. B, Heatmap of quantified immunoblot data in A and one additional replicate (Supplemental Figure 2), normalized to each control C.

D1 and CP43 were very low (D1) or not detected (CP43) in the dark, and increased progressively after 3-6h to levels similar to the continuous light control after 24-48h. Lhcbm5 was very low in the first 6h, and increased thereafter to reach levels similar to the control at 48h. PsaA and PsaC were not detected (PsaA) or very lowly accumulated (PsaC) in the dark, and increased progressively after 3-6h to levels similar to the control at 48h. PetA and PetB were already detected in the dark, and increased upon illumination to levels comparable to the control after 24-48 h. Finally, ATPb was detected in the dark to levels similar or slightly higher compared to the continuous light control, and stayed constant throughout the light treatment (Figure 4 and Supplemental Figure 2).

Together, our results show that PSI and PSII proteins rapidly accumulated after the light onset, consistent with rapid photosystem assembly under illumination, concomitant with pigment accumulation (Figures 2 and 3). *b6f* proteins, already present in the dark, further accumulated in the light, correlating with the development of thylakoid membranes (Figures 2 and 3). In contrast, ATPb was already present in the dark, suggesting ATPase activity may be essential to facilitate fast energy production required to assemble the chloroplast. Importantly, in all cases, levels at 24-48h were similar to the continuous light control (Figure 4 and Supplemental Figure 2), indicating that light induces rapid chloroplast biogenesis, in agreement with the early and progressive gain of chloroplast functionality detected after light onset (Figure 3).

## Discussion

Understanding chloroplast biogenesis has long been a key research focus given the importance of photosynthesis and chloroplast metabolism. In angiosperms, chloroplast biogenesis has been widely studied during proplastid-to-chloroplast development (in meristematic cells, monocot leaves, and cell cultures), or etioplast-to-chloroplast development (during seedling deetiolation, particularly in Arabidopsis) (Jarvis and López-Juez, 2013; Hernández-Verdeja et al., 2020; Pipitone et al., 2021). Distinct phases of chloroplast development have been described (Pipitone et al., 2021), in a process controlled by the master regulators GLKs (Hernández-Verdeja and Lundgren, 2024). However, the recent discovery of additional regulatory players in *Marchantia polymorpha* (Frangedakis et al., 2024) underscores the value of evolutionary studies in other organisms. In Chlamydomonas, chloroplast biogenesis from a de-differentiated chloroplast occurs during zygospore germination, a process challenging to study (Cardador et al., 2025). Consequently, the yellow-in-the-dark *chIL* mutant is an attractive alternative. Here, we have shown that light treatment initiates chloroplast biogenesis in dark-adapted *chIL* cells, in a process that triggers rapid accumulation of pigments, photosynthesis-related proteins, and detection of functional chloroplast activity within hours, based on starch reserves. Interestingly, GLK-like sequences are absent in Chlamydomonas (Hernández-Verdeja and Lundgren, 2024), suggesting regulation of chloroplast biogenesis by yet unknown regulators.

Given that chloroplast proteins are encoded by both the plastid and nuclear genomes, coordination of the two is necessary for chloroplast biogenesis (Jarvis and López-Juez, 2013). In the light, plastid-to-nucleus retrograde signals (RS) in plants operate during chloroplast biogenesis (biogenic RS), being the plastid-localized pentatricopeptide protein GENOMES UNCOUPLED 1 (GUN1) a key mediator (Pesaresi and Kim, 2019; León et al., 2025). Treatment with lincomycin, an inhibitor of plastid translation, has been widely used to study biogenic RS (Martín et al., 2016). Here, we have found that lincomycin specifically affects chloroplast development and physiology in Chlamydomonas, blocking pigment accumulation and photosynthesis, and affecting growth under autotrophic conditions. In dark-grown *chIL* cells transferred to light, we have shown that lincomycin prevents chloroplast biogenesis, therefore establishing lincomycin as a valuable tool for investigating RS and chloroplast biogenesis in Chlamydomonas. The absence of GUN1 in the Chlamydomonas genome (Nishiyama et al., 2018) suggests alternative biogenic RS and mediators (Duanmu et al., 2013).

In summary, our results show that the yellow-in-the-dark *ChIL* mutant, carrying a mutation in the CHLL subunit of the DPOR (Suzuki and Bauer, 1992), is a powerful model to investigate time-resolved, light-induced chloroplast biogenesis, and to explore the poorly understood regulatory mechanisms of chloroplast assembly or the nature of the biogenic RS. By providing a detailed protocol and an initial characterization of the growth, chloroplast biochemistry, and physiology of the dark-adapted *ChIL* cells upon light exposure (with and without lincomycin), we aim to facilitate broader use of this model to dissect how photosystems assemble and chloroplasts acquire functionalization. This knowledge can inform strategies to enhance photosynthesis, improve stress tolerance, and optimize bioproduct production.

## Materials and methods

### Strains, media, and growth conditions

*C. reinhardtii* strains CC-124 and CC-3269, and the *yellow-in-the-dark* mutant CC-3840 (*chIL*) (Suzuki and Bauer, 1992) were obtained from http://www.chlamycollection.org, and maintained in Tris-acetate-phosphate (TAP)-agar plates with trace elements (Kropat et al., 2011) at 25ºC in white light (30 µmol m^-2^ s^-1^). Strains were re-streaked onto fresh TAP-agar plates one week before use. Liquid cultures were grown under white light (50 µmol m^-2^ s^-1^) or in the dark in TAP or Tris-phosphate (TP) media (Harris, 2009) with shaking (100 rpm) at 25°C. Lincomycin (Sigma L6004) was used at 0.5 mM. To obtain dark-acclimated cells, colonies were inoculated in TAP liquid media and grown for 5 days in the dark, with two rounds of dilutions and media refreshment. For dark-to-light transitions, media was replaced with fresh TP on day 5 at 1-2 × 10^6^ cells mL^-1^, kept 3 hours in the dark and transferred to light (50 µmol photons m^−2^ s^−1^). Lincomycin was added 12h before transition to light in TAP cultures, and added again before exposure to light when media was replaced with TP. Continuous light *chlL* cultures were grown in TP media. Growth curves are mean OD_750_ nm values or cell number/mL of at least triplicates.

### Chlorophyll fluorescence-based measurements

*In vivo* chlorophyll fluorescence was measured at room temperature using a pulse modulated amplitude fluorimeter (DUAL-PAM-100, Heinz Waltz GmbH, Germany). Measurements were performed in TP media after 10 min of dark adaptation with stirring. Basal (F_o_) and maximum (F_m_) fluorescence signals estimate the maximum quantum yield of photosystem II (*F_v_/F_m_*). Measurements of *F*_m_ were done in red light (25 µmol photons m^−2^ s^−1^) adding 20 µM DCMU. Maximum amount of photo-oxidizable P700 (*P*_m_) corresponds to the change in absorbance at 830 nm relative to 875 nm after application of a saturating red-light pulse (5000 µmol photons m^−2^ s^−1^, 200 ms) in far-red light (75 W m^−2^) background.

### Oxygen evolution and consumption measurements

Oxygen evolution was determined using a Clark-type oxygen electrode (Chlorolab 2+ System; Hansatech, Norfolk, UK) at 25°C in the presence of 5 mM NaHCO3 under constant stirring. Cells were dark-adapted for 5 min, exposed to 50 µmol photons m^−2^ s^−1^ for 5 min, and kept in darkness for 5 min. Respiration rate was measured as oxygen consumption in the dark. Rate of photosynthetic oxygen evolution was calculated as the difference between oxygen evolution in the light and oxygen consumption in the dark, right after the light treatment. Values were normalized to cell number.

### Pigment extraction and quantification

Total chlorophylls and carotenoids were extracted in methanol and quantified following Porra et al. (Porra et al., 1989) and Lichtenthaler (Lichtenthaler, 1987), respectively. Values were normalized to cell number. To generate the chlorophyll and carotenoid absorption spectra, samples were extracted in 85% acetone buffered with 50 mM Tris-HCl (pH=8) (Chazaux et al., 2022). Protochlorophyllide and chlorophyllide were extracted and measured as in Pineau et al. (Pineau et al., 1986).

### Light Microscopy

Cells were fixed with 2.5% (v/v) glutaraldehyde (G5882; Sigma-Aldrich) in 50 mM Tris–HCl (pH 7.5) for 1 h at 4 °C, washed with 50 mM Tris–HCl (pH 7.5) and observed using a Zeiss AXIO Scope A1 optical microscope equipped with DIC optics. Images were acquired with an Axiocam 105 camera (Zeiss) and processed using Zen v.2.3 software (Zeiss).

### Electron Microscopy

Cells were fixed with 2.5% (v/v) glutaraldehyde (G5882; Sigma-Aldrich) in 0.1M sodium-cacodylate buffer (pH 7.4) for 2 h at 25°C, washed with and postfixed with the same buffer containing 1% (w/v) OsO4 for 1h at 4ºC. Fixed cells were stained with 2% uranyl acetate for 2h, dehydrated through a gradient acetone series (from 50% to 100%), and infiltrated and embedded in Spurr resin (M0300;Sigma-Aldrich). Ultrathin sections (70 nm) obtained using an ultramicrotome (Leica; UC7) were post-stained with 2% uranyl acetate followed by lead citrate and examined using a Zeiss Libra 120 transmission electron microscope with an on-axis mounted TRS camera.

### Starch extraction and quantification

Cells were treated in 80% ethanol and pellet was subjected to alkaline hydrolysis with KOH 0.2 M at 95°C for 30 min, and neutralized with 1 M acetic acid. Starch was further hydrolyzed with amyloglucosidase (Sigma-Aldrich, A7095). Levels were determined indirectly using the glucose oxidase/peroxidase method (Sigma-Aldrich, GAGO20), and normalized to cell number.

### Protein Preparation and immunobloting

Cells were extracted in lysis buffer (50 mM Tris-HCl pH 7.5, 2% SDS, 10 mM EDTA, and 1× protease inhibitor cocktail (Sigma-Aldrich; 11836153001) at 37ºC during 30 min, and quantified with the bicinchoninic acid solution (Sigma-Aldrich; B9643). Fifteen µg of protein were subjected to 12% SDS-PAGE and blotted into nitrocellulose membranes (Amersham, 10600003). Antibodies from Agrisera (Vännäs, Sweden) and dilutions used were: D1 (AS05084, 1:10000), CP43 (AS111787, 1:3000), Lhcbm5 (AS09408, 1:10000), PsaA (AS06172, 1:5000), PsaD (AS09 461, 1:3000), petA (AS06119, 1:15000), petB (AS184169, 1:7500), and ATPb (AS05085, 1:5000). An anti-rabbit horseradish peroxidase-conjugated antiserum was used for detection. Blots were developed with the Luminata Crescendo detection system (Millipore, WBLUR0100) and visualized using ImageQuant 800 (Amersham). Densitometric quantification was done with ImageJ (NIH, USA). Data was normalized to levels in control sample, and heatmap was constructed using GraphPad Prism 8 (San Diego, USA).

## Supplemental data

**Supplemental Figure 1**. Original traces of chlorophyll fluorescence and P700+ fast kinetics used to obtain F_v_/F_m_ and P_m_ levels.

**Supplemental Figure 2**. Additional Immunoblots and quantification of photosynthesis-related proteins during chloroplast biogenesis in *chIL*.

## Acknowledgements

We thank Alizée Malnoë for her suggestion to employ the *yellow-in-the-dark* mutants in our research, and Manuel J. Mallén-Ponce for his expert guidance on photosynthetic measurements and constructive scientific discussions. We thank Cristina Vaquero and Miguel Barragán at the electron microscopy service of CITIUS-US for electron microscopy (EM) sample preparation and visualization.

## Author Contributions

M.Á.R-S, M.A.DS and E.M. conceived the project and designed the experiments. M.Á.R-S, M.A.DS, M.B-T, A.M.Q-M, and E.M. planned and performed the experiments. M.Á.R-S, M.A.DS and E.M. analyzed the data. M.Á.R-S, M.A.DS, J.L.C., and E.M. wrote the manuscript. All authors approved the final version of the manuscript.

