## Supplemental Figures for "The yellow-in-the-dark *chIL* Chlamydomonas mutant as a model for time-resolved chloroplast biogenesis and physiological responses to lincomycin"

**Supplemental Figure 1**

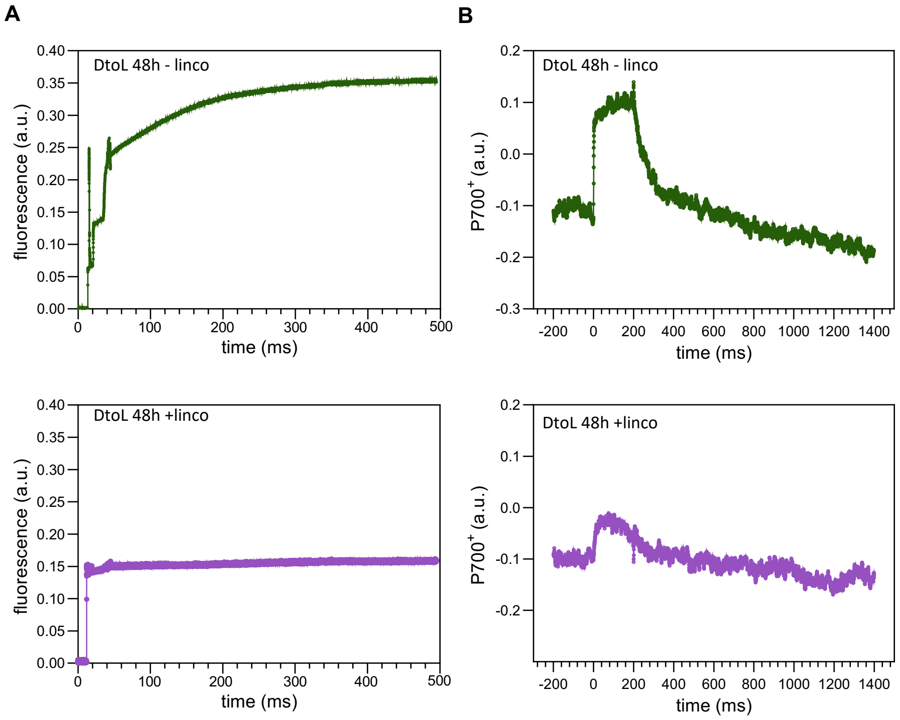


**Supplemental Figure 1**. Original traces of chlorophyll fluorescence and P700 oxido-reduction fast kinetics used to obtain Fv/Fm and Pm levels. A, Original traces of chlorophyll fluorescence kinetics used to obtain the Fv/Fm levels in Fig 3G. Data of one representative replicate of dark grown cells exposed to light during 48h without (top) and with (bottom) lincomycin. B, Original traces of P700+ fast kinetics used to obtain the Pm levels in Fig 3H. Data of one representative replicate of dark grown cells exposed to light during 48h without (top) and with (bottom) lincomycin.

**Supplemental Figure 2**

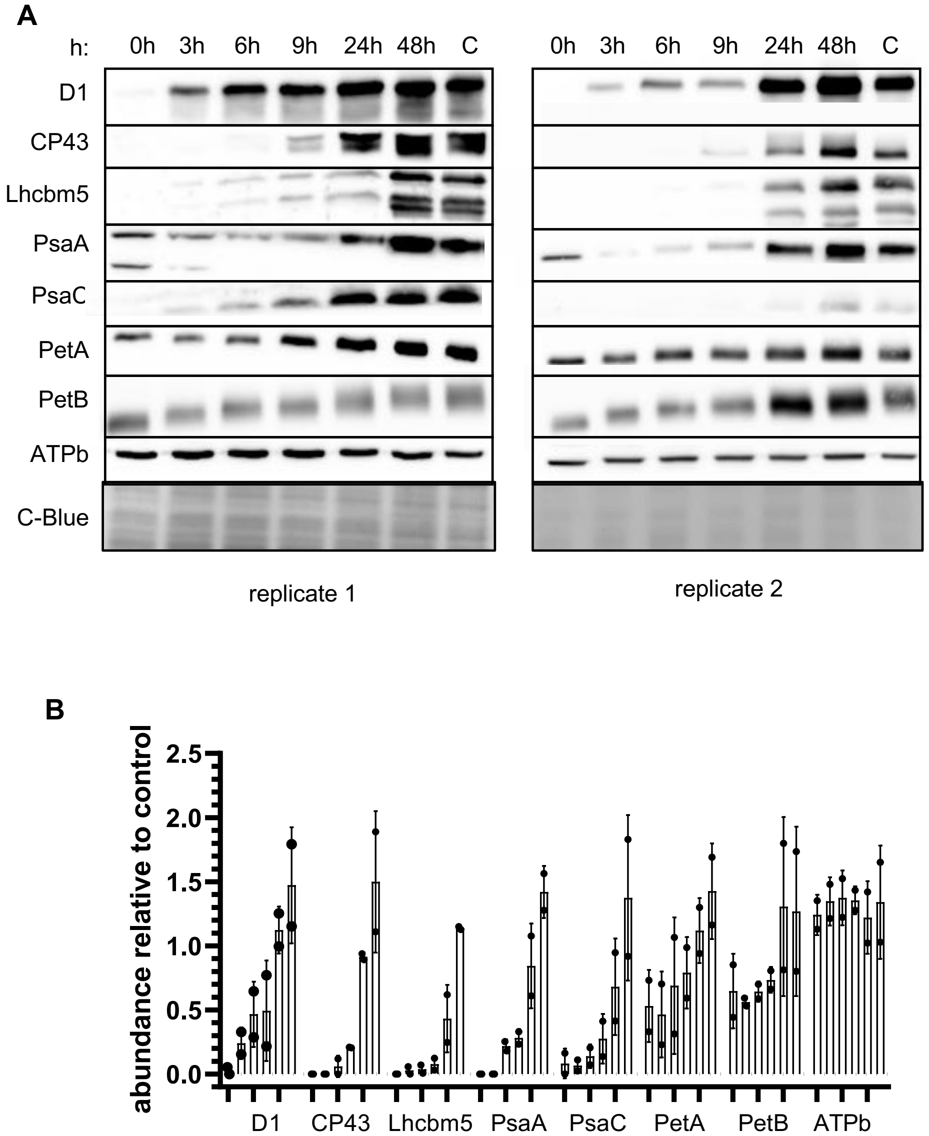


**Supplemental Figure 2**. Additional Immunoblot and quantification of photosynthesis-related proteins during chloroplast biogenesis in *ChlL*. A, Immunoblot replicates 1 (shown in Figure 4) and 2 of the accumulation of PSII (D1 and CP43), PSI (Lhcbm5, PsaA, and PsaC), Cyt $b_6f$ complex (petA and petB) and ATP synthase ATPb proteins, in dark-grown *ChlL* cells at 0, 1, 3, 6, 9, 12, 24, and 48 h after illumination. A control (C) corresponding to *ChlL* cells grown under continuous light is included. Coomassie blue (C-blue) staining served as loading control. B, Quantification of the immunoblot analysis shown in A and Figure 4A. Values are means from 2 independent experiments, and error bars represent standard deviation. Each dot represents the value in one replicate.
